# Characterisation of B.1.1.7 and Pangolin coronavirus spike provides insights on the evolutionary trajectory of SARS-CoV-2

**DOI:** 10.1101/2021.03.22.436468

**Authors:** Samuel J. Dicken, Matthew J. Murray, Lucy G. Thorne, Ann-Kathrin Reuschl, Calum Forrest, Maaroothen Ganeshalingham, Luke Muir, Mphatso D. Kalemera, Machaela Palor, Laura E. McCoy, Clare Jolly, Greg J. Towers, Matthew B. Reeves, Joe Grove

## Abstract

The recent emergence of SARS-CoV-2 variants with increased transmission, pathogenesis and immune resistance has jeopardised the global response to the COVID-19 pandemic. Determining the fundamental biology of viral variants and understanding their evolutionary trajectories will guide current mitigation measures, future genetic surveillance and vaccination strategies. Here we examine virus entry by the B.1.1.7 lineage, commonly referred to as the UK/Kent variant. Pseudovirus infection of model cell lines demonstrate that B.1.1.7 entry is enhanced relative to the Wuhan-Hu-1 reference strain, particularly under low expression of receptor ACE2. Moreover, the entry characteristics of B.1.1.7 were distinct from that of its predecessor strain containing the D614G mutation. These data suggest evolutionary tuning of spike protein function. Additionally, we found that amino acid deletions within the N-terminal domain (NTD) of spike were important for efficient entry by B.1.1.7. The NTD is a hotspot of diversity across sarbecoviruses, therefore, we further investigated this region by examining the entry of closely related CoVs. Surprisingly, Pangolin CoV spike entry was 50-100 fold enhanced relative to SARS-CoV-2; suggesting there may be evolutionary pathways by which SARS-CoV-2 may further optimise entry. Swapping the NTD between Pangolin CoV and SARS-CoV-2 demonstrates that changes in this region alone have the capacity to enhance virus entry. Thus, the NTD plays a hitherto unrecognised role in modulating spike activity, warranting further investigation and surveillance of NTD mutations.

## Introduction

We are entering a new phase of the ongoing SARS-CoV-2 pandemic. Various successful vaccine strategies are promising to alleviate disease and reduce viral transmission, offering a route to societal and economic normality. However, we have simultaneously witnessed the emergence of numerous geographically distinct SARS-CoV-2 variants that may have enhanced transmission, increased disease severity and may evade/escape immunity induced by contemporaneous natural infection and/or vaccination (1–13). As global vaccinations continue apace, the course of the pandemic will be determined by how SARS-CoV-2 navigates new and existing transmission/immunological bottlenecks. It is imperative that we understand the molecular mechanisms that permit SARS-CoV-2 fitness increases/immunological escape and consider the scope for further evolution.

At the time of writing, three variants of concern are the current focus of epidemiological, clinical and virological investigation. The lineages are commonly referred to by the country in which they were originally identified, however, we will primarily use the PANGO lineage nomenclature (14): B.1.1.7 originating from the United Kingdom, B.1.351 from South Africa, and P.1 from Brazil. Each variant lineage is defined by a constellation of mutations throughout the viral genome. Whilst many of the mutations are specific to a particular lineage there are some commonalities, suggesting convergent evolution. For example, NSP6 Δ106-108 and NSP12 P323L are apparent in each of the three variants (10).

Spike protein, in particular, has accumulated multiple mutations in each variant lineage. This is notable given its importance in transmission, its dominance in natural immunity and its widespread use as an immunogen. Similar to related viruses, SARS-CoV-2 spike protein mediates entry in a step-wise manner involving: interaction of the S1 subunit with angiotensin-converting enzyme 2 (ACE2), proteolytic processing at the S2’ cleavage site and, finally, shedding of the S1 subunit, which triggers the S2 fusion machinery (15–17). Unlike many related viruses, SARS-CoV-2 possesses an additional S1/S2 polybasic cleavage site upstream of the S2’ cleavage site, allowing spike preprocessing by intracellular proteases, most notably furin, during virion assembly and release; this has been linked to efficient human-to-human transmission (18–21). The spike mutations observed in current variants can be broadly divided into three categories based on their locations: N-terminal domain (NTD) mutations including deletions found in both B.1.1.7 and B.1.351; receptor binding domain (RBD) mutations including N501Y (B.1.1.7, B.1.351 and P.1) and E484K (B.1.351 and P.1); and stalk/S2 mutations including P681H proximal to the polybasic S1/S2 cleavage site (B.1.1.7). This wide range of spike mutations suggest that SARS-CoV-2 variants will exhibit altered entry, particle stability and/or immunoresistance. Here, we investigate virus entry by B.1.1.7 spike using pseudovirus and, through characterisation of spike from closely related viruses, consider the evolutionary pathways by which SARS-CoV-2 may increase fitness and achieve immune escape. Our study demonstrates that the entry characteristics of B.1.1.7 spike are distinct from predecessor viruses. Our work also suggests that spike NTD has an, as yet unappreciated, role in regulating spike function/stability, therefore warranting further investigation and genomic surveillance.

## Results

### B.1.1.7 spike mediates efficient virus entry

Replication-deficient lentiviruses, encoding a luciferase reporter gene, were pseudotyped with SARS-CoV-2 spike to provide a surrogate measure of SARS-CoV-2 entry. We initially compared the entry of pseudovirus (PV) bearing B.1.1.7 spike to those with spike from the Wuhan-Hu-1 reference strain (Wu-Hu-1). Western blotting of lysates from cells producing PV show equivalent expression and proteolytic processing of Wu-Hu-1 and B.1.1.7 spike (Fig. 1A, top). Parallel measurements of spike in PV pellets indicate a subtle, but consistent, reduction in B.1.1.7 spike incorporation into virus particles (Fig. 1A, bottom, also see Fig. 2B & 3C). This is similar to the B.1.1.7 spike incorporation, in PV and authentic virions, apparent in the recent work of Brown et. al. (22). We assessed PV infection of three model cell lines commonly used for SARS-CoV-2 infection; HeLa ACE2 (stably expressing exogenous ACE2), Calu-3 (with endogenous ACE2) and HEK 293T (without introduction of exogenous ACE2). Notably, despite decreased spike incorporation, B.1.1.7 PV achieved greater entry than Wu-Hu-1 PV in each cell type, and had a particularly large advantage in HEK 293T cells, where B.1.1.7 PV exhibited ~10 fold enhancement relative to Wu-Hu-1, compared to ~3 fold in HeLa ACE2 and Calu-3 (Fig. 1B & C).

**Fig. 1.**
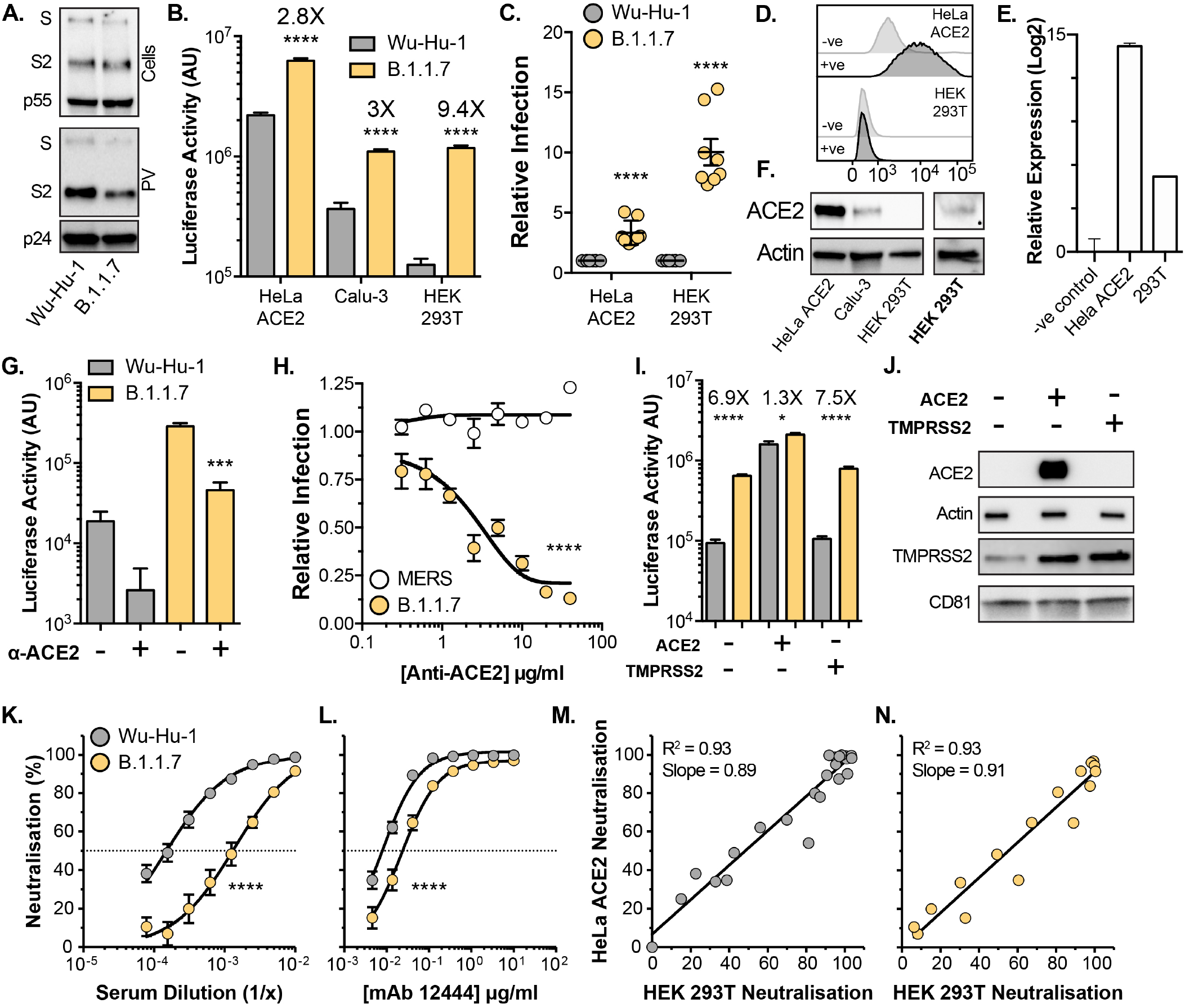
Entry characteristics of B.1.1.7 spike. Lentiviral pseudovirus (PV), encoding a luciferase reporter gene, were used to evaluate spike-mediated entry by Wu-Hu-1 (Wuhan-Hu-1 reference strain) and B.1.1.7. **A.** Cellular expression and PV incorporation of spike were assessed by western blotting using an anti-S2 mAb. Lentiviral capsid components were detected using anti-p24/55. **B.** Representative raw luciferase activity upon PV infection of the stated cell lines, annotated values represent fold enhancement of B.1.1.7 entry compared to Wu-Hu-1, n=4 technical repeats **C.** B.1.1.7 infection of HeLa ACE2 and HEK 293T cells relative to Wu-Hu-1 control. Data points represent the mean of independent experiments, n=8 **D.** Surface expression of ACE2 (goat anti-ACE2) in HeLa ACE2 and HEK 293T cells was assessed by flow cytometry, negative control represents cells incubated with secondary antibody only. **E.** ACE2 RNA transcripts were assessed by qPCR, abundance is expressed relative to negative control (no-RT). **F.** Western blot detection of ACE2 (rabbit anti-ACE2) in cell lysates of the stated cells. ACE2 was detected in HEK 293T samples after loading 3X more lysate and increasing signal exposure time (labelled in boldface). **G.** Representative raw luciferase activity after PV infection of HEK 293T cells preincubated with 20*μ* g/ml goat anti-ACE2, n=4 technical repeats **H.** B.1.1.7 and MERS-CoV PV infection of HEK 293T cells treated with a serial dilution of anti-ACE2, data points are expressed relative to untreated control cells and represent the mean of n=3 independent experiments. **I.** Raw luciferase values after Wu-Hu-1 and B.1.1.7 PV infection of HEK 293T cells transfected to over-express ACE2 or TMPRSS2, annotated values represent fold enhancement of B.1.1.7 entry compared to Wu-Hu-1, n=8 technical repeats **J.** Over-expression confirmed by western blotting, actin and CD81 were used as loading controls for ACE2 and TMPRSS2 respectively. **K.** & **L.** Neutralisation of PV infection of HeLa ACE2 cells by a reference convalescent serum (20/130) and mouse anti-RBD mAb 12444. Data points represent the mean of n=3 independent experiments. **M.** & **N.** Scatter plots demonstrating near-perfect linear correlation between neutralisation values derived from HeLa ACE2 and HEK 293T. Full neutralisation curves for either cell line provided in Fig. S1. In all plots, error bars indicate standard error of the mean, statistical analysis (T-test) performed in GraphPad Prism. Fitted curves were determined to be statistically different using an F-test (denoted by asterisks).

**Fig. 2.**
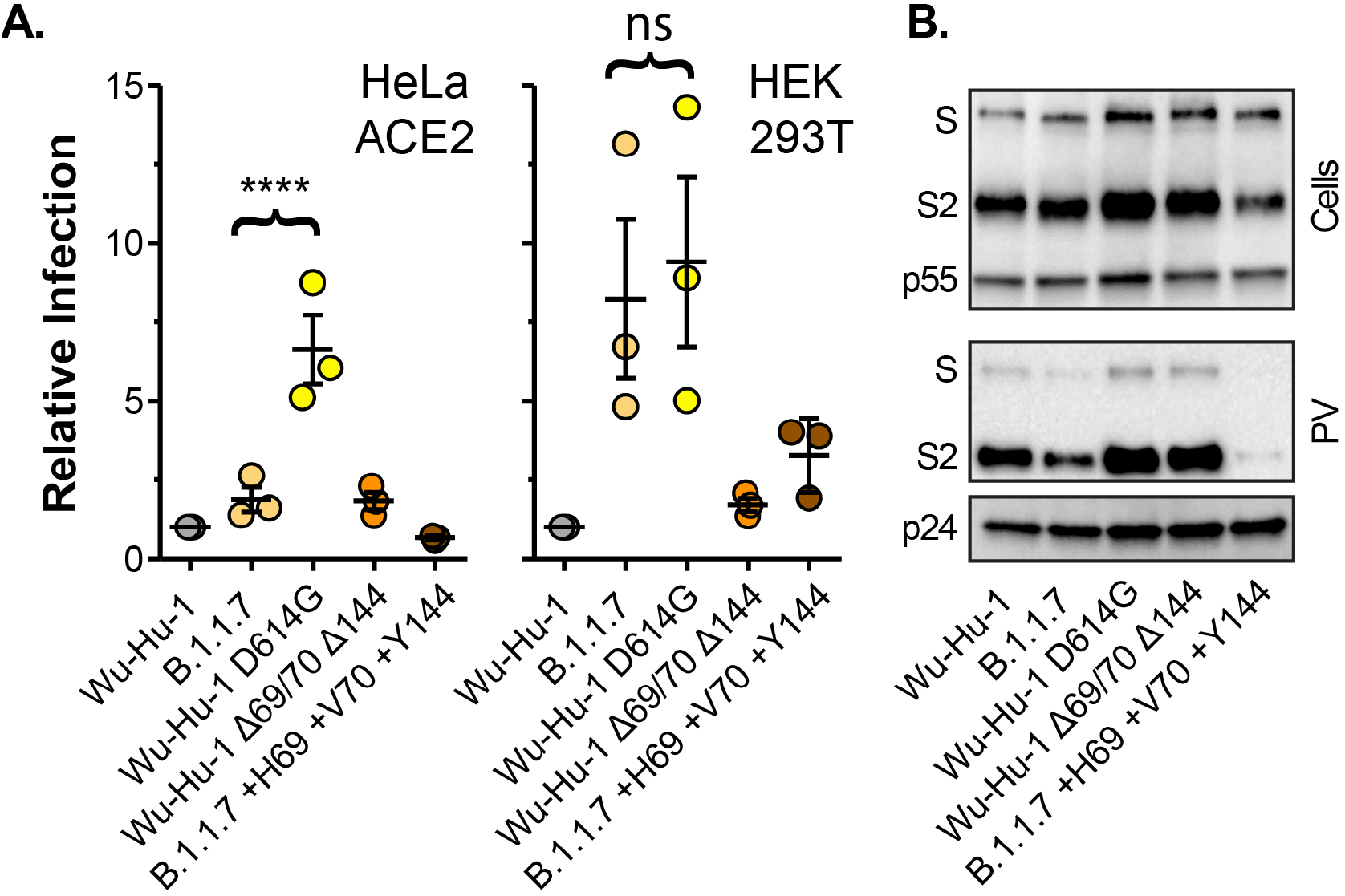
Molecular determinants of B.1.1.7 spike entry phenotype. **A.** Entry of PV bearing the stated spike proteins into HeLa ACE2 and HEK 293T. Infection is expressed relative to Wu-Hu-1, data points represent mean values from n=3 independent experiments. **B.** Cellular expression and PV incorporation of the stated spike proteins were assessed by western blotting. In all plots, error bars indicate standard error of the mean, statistical analysis (one-way ANOVA) performed in GraphPad Prism.

There are mixed reports regarding whether HEK 293T cells express the canonical SARS-CoV-2 receptor, ACE2, and alternative receptors have been proposed in this cell line (23–26). This raises the possibility that B.1.1.7 may be entering through an ACE2-independent pathway. To assess this we first examined ACE2 expression in HEK 293T cells. Whilst cell surface ACE2 was undetectable in HEK 293T cells by flow cytometry (Fig. 1D), qPCR indicated the presence of ACE2 transcripts albeit at a >500 fold lower level than HeLa ACE2 cells (Fig. 1E). ACE2 was readily observed by western blotting in HeLa ACE2 and Calu-3 cell lysates, whereas detection in HEK 293T required greater protein loading and increased signal exposure time (Fig. 1F). These data suggest that although HEK 293T endogenously express ACE2, protein levels are at the edge of detection by western blot and substantially lower than in the other cell types studied here. We then used anti-ACE2 receptor blockade to further examine the route of SARS-CoV-2 entry into HEK 293T. Entry of Wu-Hu-1 and B.1.1.7 PV into HEK 293T was inhibited by an ACE2 antibody previously shown to block SARS-CoV-2 infection (16) (Fig 1G). Inhibition was dose dependent and the antibody was ineffective against middle eastern respiratory syndrome CoV spike PV (Fig 1H), which does not use ACE2 as a receptor, thus demonstrating specificity. From these data we conclude that B.1.1.7, and Wu-Hu-1, entry into HEK 293T cells is occurring via an ACE2-dependent pathway and is unlikely to have switched to an alternative receptor.

We reasoned that the relatively high ACE2 expression levels observed in HeLa ACE2 and Calu-3 cells (Fig 1F) may compensate for the relative inefficiency of Wu-Hu-1 entry, whereas the low level ACE2 expression in HEK 293T cells revealed optimised entry by B.1.1.7 spike. To test this we examined PV infection of transfected HEK 293T over-expressing either ACE2 or the spike activating protease, TM-PRSS2 (Fig 1I). Over-expression of ACE2, but not TMPRSS2, preferentially enhanced Wu-Hu-1 spike PV infection over B.1.1.7, therefore eliminating the differential between Wu-Hu-1 and B.1.1.7 spike. This is consistent with the hypothesis that B.1.1.7 spike may be optimised for entry into poorly permissive cells, an advantage lost in cells that are highly susceptible to SARS-CoV-2. Of note, we observed endogenous expression of TMPRSS2 in HEK 293T cells (Fig 1J), and this was enhanced by over-expression of ACE2 alone suggesting interdependence in the expression/stability of these coronavirus entry factors. This may relate to the previously reported interactions between ACE2 and TMPRSS2 (27).

To further dissect entry by B.1.1.7 into HeLa ACE2 and HEK 293T cells we used spike specific antibodies in a virus neutralisation assay, reasoning that divergent entry pathways may exhibit different patterns of antibody sensitivity. We examined neutralisation of PV infection of HeLa ACE2 and HEK 293T cells by a reference convalescent serum, collected during the first wave of the pandemic (thus prior to the emergence of B.1.1.7), and two RBD-specific mouse monoclonal antibodies (Fig. 1 K & L and Fig. S1). Wu-Hu-1 spike was potently neutralised by each antibody treatment, whereas B.1.1.7 exhibited reduced neutralisation, with an 8 fold decrease in sensitivity to the reference serum (IC50 of 1/6600 vs 1/800) and complete resistance to one anti-RBD monoclonal antibody (Fig. S1; mAb 12443). This pattern of antibody sensitivity was identical when measured in either HeLa ACE2 or HEK 293T cells, as demonstrated by the near-perfect linear relationship between neutralisation values in either cell line (Fig. 1M & N). Taken together, our data suggest that the overall mode of entry mediated by B.1.1.7 spike remains unchanged relative to Wu-Hu-1, but it may allow for enhanced infection under suboptimal conditions, such as limited ACE2 expression.

### The molecular determinants of B.1.1.7 entry efficiency

Thus far we have compared B.1.1.7 to the Wuhan-Hu-1 reference strain. However, B.1.1.7, and all current variants of concern, arose from the previously dominant variant which, unlike Wu-Hu-1, contains a D614G spike mutation. D614G is associated with increased viral fitness, likely through reduced shedding of the S1 subunit and/or greater occupancy of the RBD ‘up’ conformation of spike, which may promote ACE2 interaction, but with modest effects on neutralisation sensitivity (28–33). We therefore compared Wu-Hu-1, B.1.1.7 and Wu-Hu-1 bearing the D614G mutation (Fig. 2A). Wu-Hu-1 D614G permitted greater infection of HeLa ACE2 cells than both Wu-Hu-1 and B.1.1.7 spike, whereas in HEK 293T cells Wu-Hu-1 D614G and B.1.1.7 spike exhibited equivalent activity. Again, when compared to Wu-Hu-1 and Wu-Hu-1 D614G, spike incorporation was modestly reduced in B.1.1.7 PV (Fig. 2B). These data demonstrate that, by comparison to the previous dominant variant (i.e. virus bearing the D614G mutation), the spike mutations found in B.1.1.7 have a negative effect on virus entry into some cell types (i.e. HeLa ACE2 cells) whilst maintaining efficient entry into others (HEK 293T).

Deletions within the NTD are a recurring feature in recent SARS-CoV-2 variants including B.1.1.7, which is lacking H69 V70 (Δ69/70), and Y144 (Δ144). To investigate the contribution made by these mutations, we generated Wuhan-Hu-1 spike in which these residues had been genetically deleted (Wu-Hu-1 Δ69/70 Δ144), and B.1.1.7 spike in which they were restored (B.1.1.7 +H69 +V70 +Y144). Wu-Hu-1 Δ69/70 Δ144 PV was indistinguishable from Wu-Hu-1, indicating that these NTD deletions alone are insufficient to enhance entry (Fig. 2A). Strikingly, restoring H69 V70 and Y144 abolished the B.1.1.7 entry phenotype (i.e. we observed Wu-Hu-1-like levels of PV infection of HeLa ACE2 and HEK 293T cells). This effect is likely attributable to a marked reduction in spike protein levels and PV incorporation in B.1.1.7 +H69 +V70 +Y144 (Fig. 2B). This suggests that mutations within B.1.1.7 have deleterious effects on spike protein stability that are compensated for by second site suppressor mutations, in this case the NTD deletions. Therefore the NTD has an as yet unrecognised role in regulating/rectifying protein activity and/or facilitating adaptations elsewhere in spike.

### Entry characteristics of closely related sarbe-coviruses

Of the three major subdomains in spike (NTD, RBD and S2), the NTD is the most genetically diverse across the sarbecoviruses, the family of -coronaviruses to which SARS-CoV-2 belongs (Fig. 3A and S2A). However, this diversity is not distributed equally across the NTD (Fig. S2B). Short inframe deletions, such as those in B.1.1.7, map to regions of genetic insertion/deletion (InDel) that are found across the sarbecoviruses. As defined by Garry et. al. (34, 35), there are five InDel regions in the NTD (Fig. S2B); whilst these are interspersed throughout the linear protein sequence, they map to disordered loops that are closely juxtaposed on the outward facing tip of spike (Fig S2C). Notably these loops are thought to protrude from the glycan shield that masks much of the surrounding protein surface (36). The deletions found in B.1.1.7 occur in NTD InDel regions 2 (Δ69/70) and 3 (Δ144), whereas the Δ242-244 deletion found in B.1.351 (South African variant) occurs in InDel region 5 (Fig S2D). Importantly, 90% of the S gene deletions found within the GISAID database correspond to InDel regions within the NTD, with Δ69/70 and Δ243-244 being the most frequent deletions in their respective regions, appearing across numerous independent lineages (37–39). These InDel regions also overlap with epitopes for many potently neutralising NTD-specific mAbs (40, 41), leading to the inference that the emergence of such mutations are the result of immune selection.

**Fig. 3.**
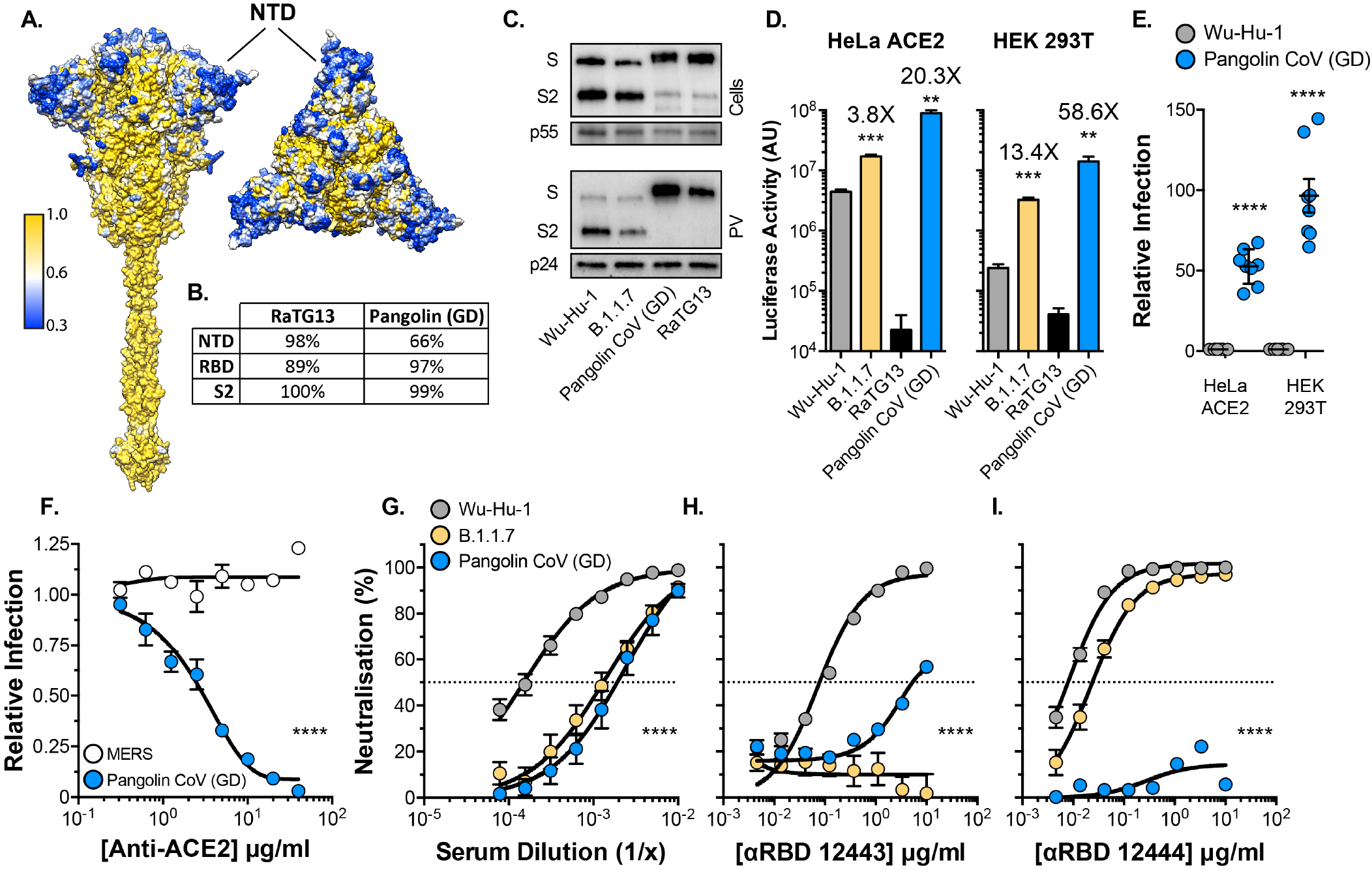
Entry characteristics of Pangolin CoV spike. Lentiviral PV encoding a luciferase reporter gene were used to evaluate spike-mediated entry by RaTG13 and Pangolin CoV (Guangdong isolate, GD). **A.** Surface representation of spike protein color coded for conservation across the sarbecoviruses, spike is shown from the side and from above. NTD, annotated, is a hotspot of diversity. **B.** Table provides percent protein homology of RaTG13 and Pangolin CoV (GD) spike domains in comparison to SARS-CoV-2. **C.** Cellular expression and PV incorporation of the stated spike proteins were assessed by western blotting. **D.** Representative raw luciferase activity upon PV infection of the HeLa ACE2 and HEK 293T cells by the stated PV, annotated values represent fold enhancement in entry compared to Wu-Hu-1, n=3 technical repeats **E.** Pangolin CoV entry into HeLa ACE2 and HEK 293T cells relative to Wu-Hu-1 control. Data points represent the mean of independent experiments, n=8. **F.** Entry of PV bearing the stated spike proteins into HEK 293T cells treated with a serial dilution of anti-ACE2, data points are expressed relative to untreated control cells and represent the mean of n=3 independent experiments. **G.-I.** Neutralisation of PV infection of HeLa ACE2 cells by a reference convalescent serum (20/130) and mouse anti-RBD mAbs 12443 and 12444. Data points represent the mean of n=3 independent experiments. In all plots, error bars indicate standard error of the mean, statistical analysis (T-test) performed in GraphPad Prism. Fitted curves were determined to be statistically different using an F-test (denoted by asterisks).

The zoonotic origin of SARS-CoV-2 remains unknown, however, very closely related viruses have been identified in bats and pangolins. For example, the spike of the bat coronavirus RaTG13 is ~97% similar to that of SARS-CoV-2 with the majority of divergence occurring within the RBD (Fig 3B). By contrast, the Pangolin CoV (Guangdong isolate) spike is more distantly related to SARS-CoV-2 (~90% similarity), but with a greater similarity within the RBD (~97%). Notably, the NTDs of Guangdong (GD) and Guangxi (GX) Pangolin CoV are the major sources of divergence from SARS-CoV-2 and possess InDel polymorphisms proximal to those found in the emergent SARS-CoV-2 variants (Fig S2D). Guangdong Pangolin CoV in particular has fewer residues than SARS-CoV-2 (Wu-Hu-1) within InDel regions 2, 3 and 5, analogous to the deletions found in B.1.1.7 and B.1.351. Therefore, we sought to examine the entry characteristics of RaTG13 and Pangolin CoV (GD) spike to gain insights on how spike diversity may influence protein function and antibody cross-neutralisation.

Spike proteins from Pangolin CoV (GD) and RaTG13 were readily expressed and incorporated into PV, albeit without prior cleavage at the S1/S2 junction due to the lack of the polybasic site found in SARS-CoV-2 (Fig. 3C). Recent structural/binding studies with these spike proteins suggest that the RBD of RaTG13 exhibits very low affinity for human ACE2 (42), indeed, we observed minimal infection by RaTG13 PV in HeLa ACE2 or HEK 293T cells (Fig. 3D). By contrast, Pangolin CoV (GD) spike has been shown to have a similar, if not slightly higher, affinity for human ACE2 compared to that of SARS CoV-2 spike (43), consistent with the high sequence homology within the RBD. Therefore, it was expected that Pangolin CoV PV may achieve comparable infection to SARS-CoV-2. Surprisingly, Pangolin CoV PV exhibited 50-100 fold greater infection of HeLa ACE2 and HEK 293T cells than SARS-CoV-2 (Fig. 3D & E). As with Wu-Hu-1 and B.1.1.7 (Fig. 1G & H), the infection by Pangolin CoV of HEK 293T cells was potently inhibited by anti-ACE2 (Fig. 3F), demonstrating canonical virus entry, albeit with much greater efficiency. Next we examined cross-neutralisation of Pangolin CoV by convalescent serum and anti-RBD mAbs. Pangolin CoV PV exhibited similar levels of resistance to the reference serum as SARS-CoV-2 B.1.1.7, possibly indicating common evasion mechanisms (Fig. 3G). Notably, Pangolin CoV PV resisted neutralisation by either anti-RBD mAbs (Fig. 3H-I), despite the high sequence similarity in the SARS-CoV-2 and Pangolin CoV RBDs.

### N-terminal domain contains determinants of spike activity

Given the importance of the NTD deletions in B.1.1.7 spike (Fig. 2) and the divergence of Pangolin CoV NTD, including deletions within InDel regions 2, 3 and 5 (Fig S2D), we hypothesised that the NTD may contribute to high-efficiency entry by Pangolin CoV. To investigate this we performed genetic swaps of the NTDs of Wu-Hu-1 and Pangolin CoV (GD) spike and examined PV infection (Fig. 4). Remarkably, despite bearing no changes to the RBD or downstream fusion machinery in S2, Wu-Hu-1/Pangolin CoV NTD phenocopied the B.1.1.7 variant, exhibiting ~3 and ~10 fold enhanced infection of HeLa ACE2 and HEK 293T cells respectively (Fig. 4B). This demonstrates that changes within the NTD alone have the capacity to increase spike activity. By contrast, the entry of Pangolin CoV/Wu-Hu-1 NTD was indistinguishable from Wu-Hu-1. This may be attributable to an apparent reduction in spike protein levels and incorporation into PV (Fig. 4C). This is strikingly similar to the phenotype of B.1.1.7 with NTD deletions H69 V70 and Y144 restored (Fig. 2), and supports the conclusion that the NTD may regulate spike stability. To summarise, the NTD contains determinants of spike protein activity and stability that likely contribute to Pangolin CoV PV’s highly efficient infection of human cells, and changes in the NTD are sufficient to enhance virus entry, in the absence of alterations in the RBD or fusion machinery.

**Fig. 4.**
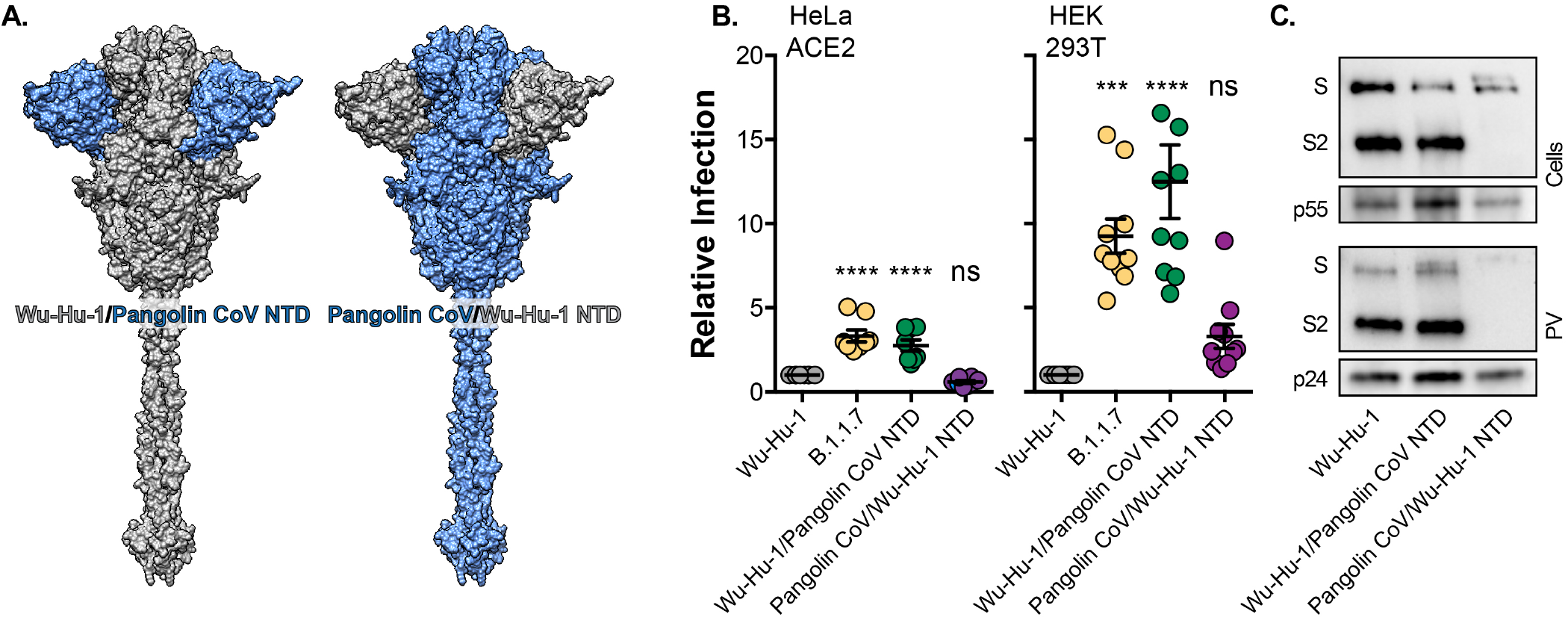
The effect of SARS-CoV-2/Pangolin CoV spike NTD swaps on virus entry. **A.** Surface representation of spike protein illustrating the NTD swaps between Wu-Hu-1 and Pangolin CoV. **B.** Entry of PV bearing the stated spike proteins into HeLa ACE2 and HEK 293T. Infection is expressed relative to Wu-Hu-1, Data points represent the mean of independent experiments, n=10. **C.** Cellular Expression and PV incorporation of the stated spike proteins were assessed by western blotting. In all plots error bars indicate standard error of the mean, statistical analysis (one-way ANOVA) performed in GraphPad Prism.

## Discussion

The recent emergence and spread of SARS-CoV-2 variants has exacerbated the ongoing public health emergency posed by the COVID-19 pandemic. Devising effective strategies to control emergent variants will require multidisciplinary investigations encompassing epidemiology, clinical science and fundamental molecular virology. The foremost questions include: in what way are the variants biologically/immunologically distinct, what selection pressures are driving SARS-CoV-2 evolution, and is there capacity for further significant change?

Our experiments demonstrate that the activity of B.1.1.7 spike is altered relative to both Wu-Hu-1 (representative of wave-one SARS-CoV-2) and Wu-Hu-1 D614G (representative of the previously dominant strain). However, we observed a complex pattern of infection. B.1.1.7 entry in HeLa ACE2 cells, which express high levels of receptor, exceeded that of Wu-Hu-1 but was lower than Wu-Hu-1 D614G. This is broadly consistent with other reports suggesting authentic SARS-CoV-2 B.1.1.7 isolates may have a replicative disadvantage in some cell types (22). By contrast, Wu-Hu-1 D614G and B.1.1.7 infection of HEK 293T cells were indistinguishable, both being ~10 fold greater than Wu-Hu-1. HEK 293T cells endogenously express very low levels of ACE2. Therefore, our data suggest that the D614G mutation represents adaptation to host and permits infection of poorly permissive cell types. Critically, this ability is maintained in B.1.1.7 (albeit through a complex set of additional mutations). Taken together, these data do not offer a simple explanation for why B.1.1.7 exhibits increased transmissibility and pathogenesis (5, 12) compared to its direct predecessor (i.e. viruses bearing D614G). Nonetheless, the pattern of infection in model cell lines by B.1.1.7 PV suggests evolutionary tuning of spike activity, which may increase entry efficiency and alter or broaden cellular tropism.

B.1.1.7 PV exhibited some antibody neutralisation resistance relative to Wuhan-Hu-1. However, we used a limited panel of antibodies and we note that many other reports demonstrate that B.1.1.7 does not escape neutralisation by the majority of COVID-19 patient serum or human mAbs (2, 11, 22, 44–46). The 8-fold resistance to a convalescent serum we observed is likely to be atypical and may indicate a prevalence of anti-NTD responses in this sample, to which B.1.1.7 is particularly resistant (7, 40, 47), discussed below. Therefore, despite our data, B.1.1.7 is unlikely to achieve significant antibody escape in vaccinated/previously infected individuals.

Pangolin CoV spike has been demonstrated to have very similar affinity for human ACE2 as SARS-CoV-2 (43, 48). Given this, we were surprised to find that entry by Pangolin CoV spike was 50-100 fold enhanced relative to Wu-Hu-1 SARS-CoV-2. This observation is important for two reasons. It provides yet another demonstration that some animal sarbecoviruses require little/no adaptation to achieve efficient entry into human cells; they are pre-adapted (49) and ‘oven-ready’ for zoonotic spillover. More pressingly, it implies that there may be evolutionary scope for SARS-CoV-2 to optimise spike and reap further gains in entry efficiency. In addition, Pangolin CoV was largely resistant to neutralisation by anti-SARS-CoV-2 antibodies, despite high sequence homology. This indicates that achieving high levels of SARS-CoV-2 seropositivity in the human population (through natural infection or vaccination) will not necessarily prevent future spillover of related viruses.

Spike NTD is a hotspot of diversity within sarbecovirus genomes (20), suggesting relatively low functional constraints on this domain. Spike glycosylation shields much of the NTD from immune recognition. Consequently, it is under-represented in the humoral immune response, which is largely focused on the RBD (50, 51). Nonetheless, a minority component of the antibody response targets an exposed NTD antigenic supersite (40, 52), corresponding to the disordered loops encoded by InDel regions 1-5 (Fig. S2). Therefore, NTD diversity may be driven by immune selection. Indeed, it has been demonstrated that SARS-CoV-2 spike deletions, such as Δ144 found in B.1.1.7, can prevent the binding and neutralisation by NTD supersite antibodies (37, 40, 47). Moreover, the NTD has recently been demonstrated to bind the haem metabolites biliverdin and bilirubin within an intramolecular binding pocket, resulting in conformational changes that confer anti-NTD antibody resistance (53).

In addition to the capacity for NTD deletions to mediate antibody escape, our work demonstrates a hitherto unappreciated role for the NTD in regulating spike function. Restoration of deleted NTD residues in the B.1.1.7 background reduced virus entry to a level similar to that of Wu-Hu-1; this was likely the consequence of a reduction in cellular/PV protein levels, suggesting spike instability. This is consistent with recent reports on viruses containing the Δ69/70 mutation (38, 39). Swapping the Pangolin CoV NTD into Wuhan-Hu-1 spike enhanced entry without altering protein levels, suggesting increased spike activity. Whilst swapping the Wuhan-Hu-1 NTD into Pangolin CoV spike diminished virus entry; again, this was correlated with a reduction in protein levels. Taken together these data indicate that changes within the NTD alone are capable of increasing virus entry, and that NTD deletions may be necessary for rectifying deleterious effects arising from mutations elsewhere in spike.

It is notable that the development of NTD deletions and, more broadly, the emergence of SARS-CoV-2 variants have been linked to persistent infection of immunodeficient patients with weak/no neutralising antibody responses (38, 54). Many immunocompromised COVID-19 patients received convalescent plasma (CP) treatment, which would provide a degree of immune selection. However, mutations/deletions were observed in some individuals without prior CP treatment and with no endogenous neutralisation activity (55, 56), arguing immune evasion is not the sole driver of SARS-CoV-2 variation. Moreover, similar/identical NTD and RBD mutations to those observed in current SARS-CoV-2 variants have emerged upon prolonged replication in cell culture which lacks any adaptive immune selection (57). Therefore, protracted infection in immunocompromised individuals may offer an evolutionary proving ground for SARS-CoV-2 to adapt to the human host through stepwise accumulation of multiple cooperative mutations throughout spike.

Important questions remain about where SARS-CoV-2 currently resides on the fitness landscape. In this regard, the highly efficient entry by Pangolin CoV spike into human cells (10-30 fold higher than B.1.1.7) sets a precedent for additional optimisation of spike function. Understanding the evolutionary pathways that guide contemporary viral variants towards peak fitness in humans, and the conditions that promote these adaptations, will inform future mitigation strategies. Thus far, studies have focussed on the contribution made by RBD mutations, and with good reason; N501Y and E484K have been implicated in increased ACE2 affinity and escape from anti-RBD responses (2, 46, 58–62). However, our study demonstrates the capacity of the NTD to modulate spike activity. We may expect further adaptation in this region and advise ongoing genetic surveillance and investigation of NTD mutations.

## Materials and methods

### Cell culture

HeLa ACE2 (a kind gift from Dr. James Voss, SCRIPPS (51)), HEK 293T cells and Calu-3 cells were maintained at 37°C in Dulbecco’s Modified Eagle Medium supplemented with 10% foetal calf serum (FCS), 1% non-essential amino acids and 1% penicillin/streptomycin.

### Plasmids

Codon optimised open reading frames encoding spike proteins were synthesised (GeneArt, Thermo Fisher) and cloned into pCDNA3.1 and/or pD603 (ATUM) expression plasmids. We note that codon optimisation offers an additional level of biosecurity as it precludes the opportunity for homologous recombination between plasmid-derived mRNA and SARS-CoV-2 upon accidental infection of cell culture. Whilst this occurrence is highly unlikely, we would recommend codon optimisation as a precaution for investigations in which spike protein mutants are created. The following sequences were generated: SARS-CoV-2 Wuhan Hu-1 (YP_009724390.1), Wuhan Hu-1 D614G, B.1.1.7 (based on Wuhan-Hu-1 coding sequence but with mutations as listed in (10)), RaTG13 CoV (QHR63300.2), Pangolin CoV Guangdong (EPI_ISL_410721), middle eastern respiratory syndrome CoV (YP_009047204.1). For deletions/insertion of NTD residues (Fig 2) we took advantage of flanking SpeI + EcoNI restriction sites to swap the NTD coding sequence between Wuhan-Hu-1 and B.1.1.7 plasmids. For Wu-Hu-1/Pangolin CoV N-terminal domain swaps (Fig. 4) we took advantage of common SpeI and BbsI restriction sites, found at identical locations on both expression plasmids, to exchange the coding sequence for residues 1-307 (encompassing all of the NTD and 16 residues of the linker region that precedes the RBD). Human ACE2 and TMPRSS2, encoded in pCAGGS, were a kind gift from Dr Edward Wright (University of Sussex).

### Antibodies

The following antibodies were used in this study: mouse anti-spike S2 (1A9, Gene Tex), mouse anti-p55/24 (ARP366, Centre for AIDS Reagents), goat anti-ACE2 (AF933, R& D Systems), rabbit anti-ACE2 (EPR4435, abcam), rabbit anti-TMPRSS2 (EPR3861, abcam), mouse anti--actin (ab49900, abcam), mouse anti-CD81 (2.131, a kind gift from Prof. Jane McKeating, University of Oxford), convalescent serum 20/130 (National Institute for Biological Standards and Control), mouse anti-RBD 12443 and 12444 (Native Antigen Company).

### Pseudovirus and entry assays

Lentiviral pseudovirus were generated as previously reported (63). HEK 293T CD81 knockout cells (which produce higher titre PV (63)), seeded in a 6 well plate, were Fugene 6 transfected with 1.3*μ*g 8.91 lentiviral packaging plasmid, 1.3*μ*g CSFLW luciferase reporter construct and 200ng of spike expression plasmid. The cells were PBS washed and refed with fresh media 24 hours post transfection. Media containing PV were harvested at 48 and 72 hours. All supernatants were passed through a 0.45*μ*m filter prior to use. PV were stored at room temperature and routinely used within 48 hours of harvest. Producer cell lysates were harvested in Laemmli buffer to confirm protein expression by western blot.

To evaluate virus entry, 50,000 cells (in 50l of media) were seeded into each well of a 96 well plate, containing an equal volume of diluted PV. Antibody pretreatment of cells/virus were performed in plate at 37°C for 1 hour prior to infection. After 72 hours, infections were readout on a GloMax luminometer using the Bright-Glo assay system (Promega). PV without spike glycoproteins (no envelope control) were used to determine the noise threshold of the assay. PV preparations, within a given batch, were not routinely matched for infectious dose; however, normalisation of test batches by genome copy number, SYBR-green product-enhanced RT (SG-PERT, (64)) or by p24 protein levels (as evidence in western blot data) did not alter our findings.

To pellet particles, 8ml of PV media was laid over a 4ml sorbitol cushion (20% D-sorbitol, 50 mM Tris, pH 7.4, 1 mM MgCl2) and concentrated by centrifugation at 120,000xg for 2 hours at 4°C. Virus pellets were harvested in Laemmli buffer for western blotting.

### PAGE and western blotting

Samples underwent SDS-PAGE in 4-20% Mini-PROTEAN TGX precast gels (Bio Rad) under reducing conditions (Laemmli Buffer and pretreatment at 95°C for 5mins) apart from TMPRSS2/CD81 gel (non-reducing SDS sample buffer). Proteins were transferred to nitrocellulose membranes, blocked in PBS +2% milk solution + 0.1% Tween-20 and then probed by overnight incubation at 4°C with the stated antibodies (diluted in blocking buffer) followed by 1 hour at RT with secondary antibodies conjugated to horseradish peroxidase. Chemiluminescence signal was measured using a Chemidoc MP (Bio Rad).

### Flow cytometry

Surface expression of ACE2 in HeLa-ACE2 and HEK 293T cells was assessed using goat polyclonal anti-human ACE2 and secondary donkey anti-Goat IgG H+L Alexa Fluor 488. Dead cells were excluded using a fixable viability dye. Data were acquired using an LSR Fortessa II (BD Biosciences) and analysed using FlowJo version 10.5.3 (FlowJo LLC, Becton Dickinson).

### Quantitative PCR (qPCR)

RNA was extracted from 250,000 cells using RNeasy Mini Kit (Qiagen). RNA quantity and quality were assessed using a NanoDrop Lite spectrophotometer (Thermo Scientific). cDNA was synthesised from 1g total RNA using the QuantiTect Reverse Transcription Kit (Qiagen) following the manufacturer’s instructions. Duplicate aliquots of each sample were processed in parallel with and without the addition of reverse transcriptase, therefore generating matched cDNA and no-RT negative controls. qPCR was performed using PowerUp SYBR Green (Applied Biosystems) using a Quantstudio 3 Real Time PCR System (Applied Biosystems). Cycling conditions were 60°C for 2 minutes, 95°C for 2 minutes followed by 40 repetitions of 95°C for 15 seconds and 60°C for 60 seconds. Data was analysed by ΔΔCt method. ACE2 primer sequences were as reported elsewhere (65), forward: AAA CAT ACT GTG ACC CCG CAT, reverse: CCA AGC CTC AGC ATA TTG AAC A. 18S ribosomal RNA control primers, forward: GTA ACC CGT TGA ACC CCA, reverse: CCA TCC AAT CGG TAG TAG CG.

### Bioinformatics

Protein sequence alignments were performed in UGENE (66) using the multiple sequence comparison by log-expectation (MUSCLE) method (67). Sequences were aligned using Wuhan-Hu-1 (YP_009724390.1) as the reference. We analysed 27 sarbecovirus spike sequences, including those used by Garry et. al. and Holmes et. al. to examine frequent sites of insertion/deletion (34, 35): QHR63300.2, QQM18864.1, EPI_ISL_412977, QIA48623.1, EPI_ISL_410721, AHX37569.1, AAP30030.1, QJE50589.1, AGZ48828.1, BCG66627.1, AVP78031.1, AIA62277.1, EPI_ISL_852605, ABD75323.1, AKZ19076.1, AIA62330.1, QDF43815.1, AIA62310.1, AGC74165.1, QDF43835.1, AGC74176.1, QDF43830.1, Q0Q475.1, Q3I5J5.1, AIA62320.1, AID16716.1, YP_003858584.1.

### Structural modelling

Molecular graphics and analysis performed with UCSF Chimera, developed by the Resource for Biocomputing, Visualization, and Informatics at the University of California, San Francisco, with support from NIHP41-GM103311 (68).

### Statistics

All statistical analysis (T-test, one-way ANOVA with Dunnet’s correction for multiple comparisons, curve fitting and F-test) were performed in GraphPad Prism. Significance denoted by asterisks: ∗ P ≤ 0.05, ∗ ∗ P ≤ 0.01, ∗ ∗ ∗ P ≤ 0.001.

## ACKNOWLEDGEMENTS

JG is supported by a Sir Henry Dale Fellowship from the Wellcome Trust and Royal Society (107653/Z/15/Z). MBR is supported by the Medical Research Council (MR/R021384/1). CJ is supported by a Wellcome Trust Investigator Award (108079/Z/15/Z). GJT is supported by a Wellcome Trust Investigator Award (220863) and Senior Research Fellowship (108183), and grants from the Rosetrees Foundation, Medical Research Council and UKRI Genotype to Phenotype Consortium. LEM is supported by a Medical Research Council Career Development Award (MR/R008698/1). This study was supported by the UCL Coronavirus Response Fund made possible through generous donations from UCL’s supporters, alumni and friends.

## AUTHOR CONTRIBUTIONS

**Conceptualisation**, J.G.; **Methodology**, J.G., M.J.M, C.F; **Investigation**, S.J.D, M.J.M, L.G.T, A-K.R, C.F, M.G, L.M, M.D.K, M.P; **Resources**, L.E.M; **Writing**, J.G. **Funding Acquisition**, J.G. L.E.M, C.J, G.J.T, M.B.R; **Supervision** J.G. L.E.M, C.J, G.J.T, M.B.R

---

**Figure S1.**
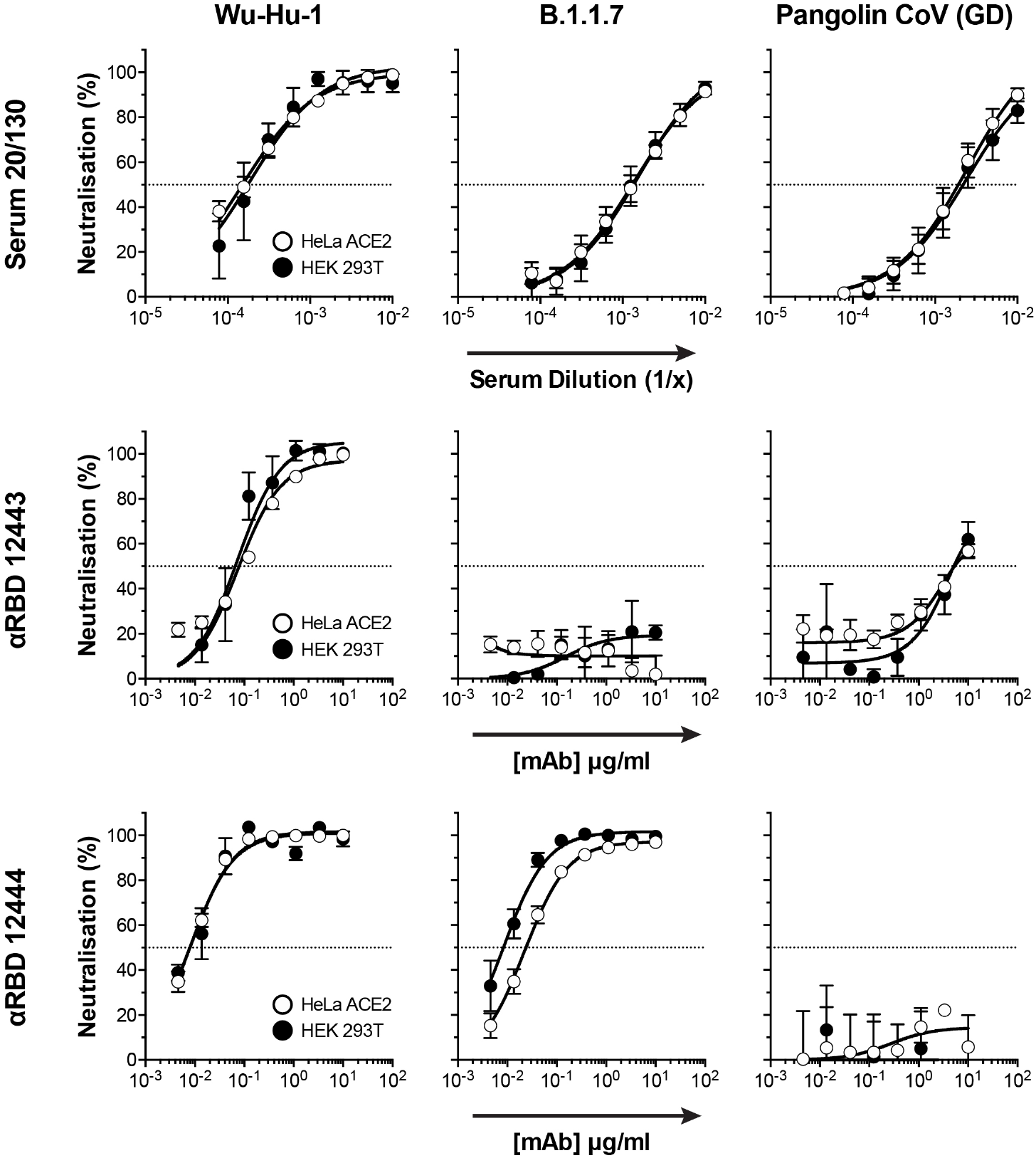
Antibody neutralisation of PV entry into HeLa ACE2 and HEK 293T cells. Neutralisation of the stated PV by a reference convalescent serum (20/130) and mouse anti-RBD mAbs 12443 and 12444. Each plot provides neutralisation values from HeLa ACE2 and HEK 293T cells. Data points represent the mean of n=3 independent experiments. Error bars indicate standard error of the mean. Curve fitting performed in GraphPad Prism, the curves on each plot (i.e. HeLa ACE2 and HEK 293T) were determined to be statistically indistinguishable.

**Figure S2.**
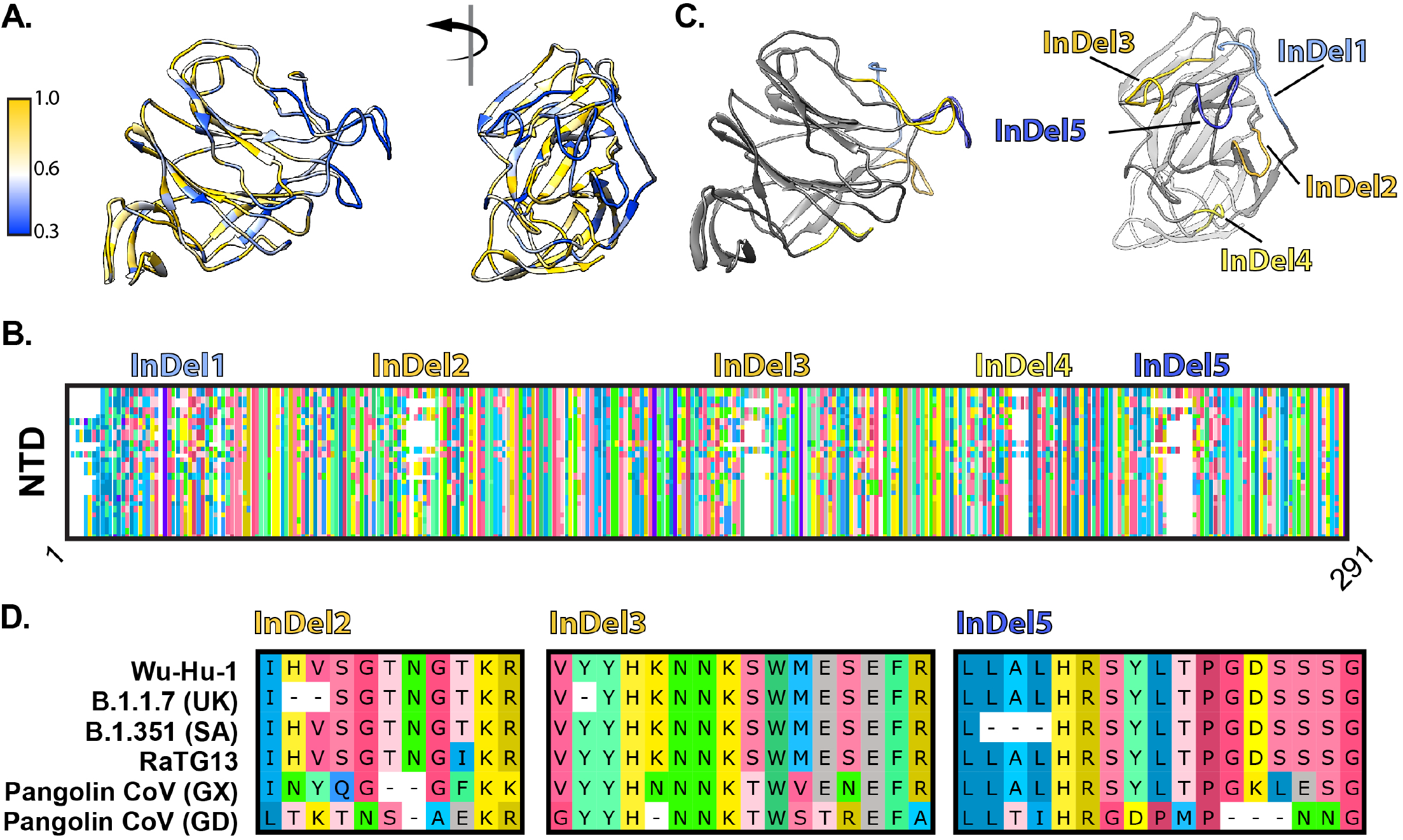
Sarbecovirus spike N-terminal domain contains InDel regions map to outwardly facing loops. **A.** Cartoon representation of a single N-terminal domain, shown from the side and en face, color coded for conservation, as denoted on the scale. **B.** Overview protein alignment of the spike N-terminal domain (residues 1-291 of SARS-CoV-2) from 28 diverse sarbecoviruses, residues color coded by amino acid identity. Annotations show the location of five regions of insertion/deletion (InDel), as defined by Garry et. al.. **D.** InDel regions map to outwardly facing loops on the NTD. **E.** Protein alignments demonstrating that current SARS-CoV-2 variants of concern and, closely related, Pangolin CoVs have deletions within InDel regions 2,3 and 5.

